# Restorative macrophage-derived RNAseT2 stimulates muscle stem cell fusion via an SLK/N-WASP/actin bundling dependent axis

**DOI:** 10.1101/2024.09.11.612435

**Authors:** Michèle Weiss-Gayet, Gaëtan Juban, Emmeran Le Moal, Antonio Moretta, Camilla Farnetani, Christelle Gobet, Jules Guillemaud, Marie-Catherine Le Bihan, Oded Shoseyov, Annie Adrait, Katharina Ternka, Odile Boespflug-Tanguy, Matthias Kettwig, Yohann Couté, Rémi Mounier, Francesco Acquati, Robert D Knight, Bénédicte Chazaud

## Abstract

Muscle stem cells (MuSCs) fuse to form myofibers to repair skeletal muscle after injury. Within the regenerative MuSC niche, restorative macrophages stimulate MuSC fusion, although the molecular mechanisms involved are largely unknown. Here, we show that restorative macrophages secrete ribonuclease T2 (RNAseT2) to stimulate MuSC fusion. RNAseT2 entered MuSCs via the mannose receptor and induced the formation of actin bundles in MuSCs, enabling cell/cell fusion. Mechanistically, RNAseT2 bound to Ste20-like kinase (SLK), which itself triggered the phosphorylation-mediated activation of N-WASP, through Paxillin phosphorylation, allowing actin bundling necessary for MuSC fusion. *In vivo*, overexpressing RNAseT2 in regenerating muscle increased fusion in newly formed myofibers in mouse and zebrafish while macrophages deficient for *RNAseT2* gene led to fusion defect and smaller myofibers. This study reveals a new function for the highly conserved RNAseT2 and provides a new molecular mechanism by which restorative macrophages support MuSC fusion during muscle repair.

## Introduction

Muscle stem cells (MuSCs) fuse to form multinucleated cells to repair skeletal muscle after a lesion ^1^. Adult MuSCs express a cell autonomous myogenesis program, but experimental manipulations *in vivo* have revealed that the environment is required for efficient muscle regeneration ^1,2^. Macrophages support myogenesis and muscle regeneration by promoting MuSC proliferation through inducing a pro-inflammatory environment then transitioning to resolution and promoting an anti-inflammatory/restorative inflammatory profile which stimulates MuSC differentiation and fusion ^2^. Although a number of molecular effectors from macrophages have been shown to regulate the various steps of myogenesis ^3^, the molecular mechanisms by which restorative macrophages stimulate MuSC fusion are still largely unknown.

Using human cells, we have previously shown that culturing MuSCs with conditioned medium from activated macrophages promotes the differentiation and fusion of MuSCs ^4^. This suggests that the main effects of anti-inflammatory macrophages on the terminal steps of myogenesis are triggered by secreted factors. To identify such factors, we performed a screen for molecules that are differentially secreted by pro-and anti-inflammatory human macrophages *in vitro* and can affect MuSC differentiation and fusion.

In this study, we show that ribonuclease T2 (RNAseT2) is expressed in restorative macrophages *in vitro* and anti-inflammatory macrophages *in vivo* in regenerating mouse muscle. RNAseT2 secreted by anti-inflammatory macrophages specifically stimulated MuSC fusion without affecting proliferation and differentiation. RNAseT2 is taken up by MuSCs via the mannose receptor and stimulates fusion through stimulation of actin remodeling and promoting increased actin bundling. We identified Ste20-like kinase (SLK) as a direct partner of RNAseT2 that triggered the phosphorylation and activation of N-WASP, a major effector of actin bundling and necessary for MuSC fusion ^5^. The ability of RNAseT2 to regulate MuSC fusion was assessed *in vivo* in mouse and zebrafish models of skeletal muscle regeneration and was shown to induce elevated fusion. In human, macrophages derived from patients carrying a mutation that results in the absence of RNAseT2 were not able to stimulate MuSC fusion as compared with normal macrophages.

## Results

### RNAseT2 is expressed by restorative macrophages and boosts muscle stem cell fusion

A differential transcriptomic analysis was performed using human macrophages activated *in vitro* into pro-inflammatory (LPS treated), alternatively activated (IL-4 treated) or anti-inflammatory macrophages (IL-10 and dexamethasone treated) and filtered for those genes encoding secreted proteins (Fig.S1A, Table S1). We previously showed that these macrophages exert differential effects on human MuSC proliferation, differentiation and fusion, with alternatively activated and anti-inflammatory macrophages stimulating their fusion ^4,6^. Among the molecules expressed strongly in anti-inflammatory macrophages and predicted to be secreted (Cluster 6 in Table 1) was Ribonuclease T2 (RNAseT2). Increased expression of RNAseT2 in anti-inflammatory macrophages relative to pro-inflammatory macrophages was confirmed by RTqPCR (Fig.S1B).

*In vivo*, analysis of inflammatory (Ly6C^pos^) and restorative (Ly6C^neg^) macrophages present in regenerating skeletal muscle 4 days after injury indicated that RNAseT2 was mainly expressed by restorative macrophages (Fig.1A). Immunostaining of regenerating mouse muscle at 8 days post-injury, a time point when only restorative macrophages are present ^7^, showed that RNAseT2 expressing cells were all positive for the pan-macrophage marker F4/80 (Fig.1B). These data indicate that restorative macrophages are the main producers of RNAseT2 *in vivo*, at the time of MuSC differentiation and fusion ^8,9^. RNAse T2 was previously identified in the human MuSC secretome during differentiation *in vitro* ^10^. However, its secretion was minimal, representing only 0.1% of the total secretome, and was secreted at lower levels during differentiation (data not shown).

**Figure 1.**
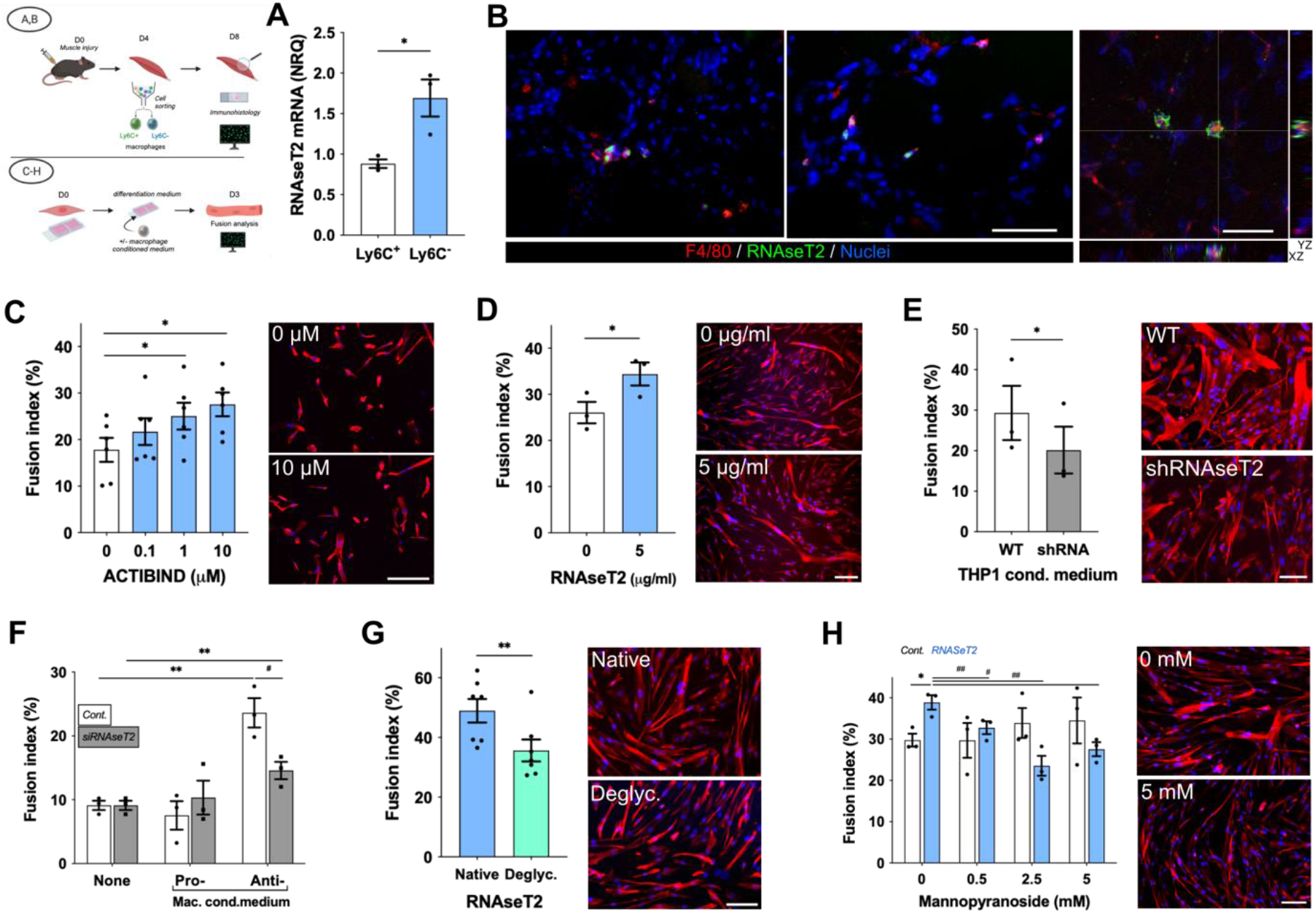
Macrophage-derived RNAseT2 stimulates human MuSC fusion. **(A)** Expression of RNAseT2 by RT-qPCR in pro-(Ly6C^pos^) and anti-(Ly6C^neg^) inflammatory macrophages in regenerating mouse muscle (4 days after a cardiotoxin-induced muscle injury). **(B)** Immunostaining for macrophages (F4/80, red) and RNAseT2 (green) (blue = Hoechst) in regenerating skeletal muscle (8 days post-injury). The right panel shows confocal microscopy colocalization on xy, xz and yz projections. **(C-H)** Fusion assay of human MuSCs after immunostaining for desmin (red) and nuclei labeling (Hoechst) of: **(C)** primary MuSCs cultured with or without ACTIBIND; **(D)** MuSCs cultured with or without recRNAseT2; **(E)** MuSCs cultured with conditioned medium from WT THP1 or shRNA-RNAseT2 THP1 anti-inflammatory macrophages; **(F)** primary MuSCs cultured with conditioned medium from pro- or anti-inflammatory primary macrophages previously treated with or without a RNAseT2-siRNA; **(G)** MuSCs cultured with native or deglycosylated RNAseT2 (5 µg/ml); (**G**) MuSCs cultured with or without RNAseT2 (5 µg/ml) and with increasing concentrations of allyl-α-D-mannopyranoside, a ligand of the mannose receptor. Results are means ± SEM of at least 3 independent experiments (*p<0.05, **, ^##^ p<0.01 using 1-way ANOVA (B), 2-ways ANOVA (E,H) or paired t-test (C,D,G)). Bars = 50 µm (left and middle panels in A), 20 µm (right panel in A) and 100 µm (C-H).

RNAseT2 belongs to the transferase-type Rh/T2/s Ribonuclease family and is highly conserved among eukaryotic and prokaryotic species ^11^. It has 2 catalytic sites and 3 N-glycosylation sites ^11^. To investigate the role of RNAseT2, we first used a fungal ortholog of RNAseT2, Actibind, extracted from *Aspergillus niger* ^12^. Actibind protein was applied to human primary MuSCs cultured *in vitro*. Actibind had no impact on primary MuSC proliferation (ki67 labeling) (Fig.S1C) or differentiation (myogenin expression) (Fig.S1D), but it stimulated primary MuSC fusion in a dose-dependent manner (Fig.1C). Similar results were obtained with human immortalized MuSCs ^13^ when exposed to recombinant human RNAseT2, with fusion increased by 132% (Fig.1D) but had no impact on proliferation (Fig.S1E). To demonstrate that RNAseT2 derived from macrophages affected cell fusion, we first treated MuSCs with conditioned medium from the macrophage cell line THP1 and from an established RNAseT2 deficient THP1 cell line ^14^. MuSC fusion was decreased by 31% when macrophages were deficient for RNAseT2 (Fig.1E). We also inhibited RNAseT2 function using siRNAs (which reduced RNAseT2 expression by 74%, Fig.S1F) on human primary macrophages, activated to assume a pro- or anti-inflammatory state. As expected, conditioned medium from pro-inflammatory macrophages did not stimulate primary MuSC differentiation (Fig.S1G) nor fusion (Fig.1F) and silencing RNAseT2 had no effect. In contrast, conditioned medium from anti-inflammatory macrophages stimulated primary MuSC fusion (+260%), in line with previous studies ^4,6^. Silencing RNAseT2 in anti-inflammatory macrophages abolished their pro-fusogenic effect by 62% (Fig.1F). We did not observe any effect of silencing RNAseT2 on the capacities of anti-inflammatory macrophage to stimulate primary MuSC differentiation (Fig.S1G).

To confirm the physiological relevance of macrophage-derived RNAseT2 for MuSC fusion, we compared the expression of RNAseT2 and fusogenic properties of Ly6C^neg^ macrophages in various conditions of muscle regeneration in mouse. Ly6C^neg^ macrophages were isolated from day 4 post-injury regenerating muscle where these restorative macorphages are known to promote MuSC fusion, from fib-mdx muscle where chronic inflammation is established and macrophages exihibit a pro-inflammatory profile, and from muscle of fib-mdx mice treated with metformin, which promotes the shift of macrophages towards a restorative profile, regenerative inflammation and improves tissue repair ^15^. Macrophages from fib-mdx mice showed both reduced RNAseT2 expresion and fusogenic capacity as compared with macrophages isolated from regenerating muscle (Fig.S1H) and the two parameters were rescued by metformin treatment (Fig.S1H). These results reveal that RNAseT2 released by anti-inflammatory macrophages has a specific stimulatory effect on MuSC fusion.

### RNAseT2 enters MuSCs via the mannose receptor

Omega-1, an orthologue of RNAseT2 from *Schistosoma mansoni*, is the major component of the parasite’s eggs and is responsible for Th2 polarization of dendritic cells through the mannose receptor (CD206) expressed at their surface ^16^. In mouse MuSCs, the mannose receptor is increasingly expressed during differentiation and is involved in their fusion ^17^. We note that human primary MuSCs expressed the mannose receptor (hMRC1) during their differentiation (Fig.S1I,J). To block recognition of RNAseT2 by the mannose receptor, we treated RNAseT2 with Peptide:N-Glycosidase F, resulting in a partial deglycosylation of the protein (shown in Fig.S1K). Deglycosylated RNAseT2 was 27% less efficient than the native protein to promote MuSC fusion (Fig.1G). Then, we performed competition experiments in which allyl-α-D-mannopyranoside, a specific mannose receptor ligand, was added at increasing concentrations to the MuSC cultures. While mannopyranoside had no impact on untreated MuSCs, it blunted the stimulating effect of RNAseT2 on their fusion (Fig.1H).

These results show that RNAseT2 secreted from anti-inflammatory macrophages can enter MuSCs via the mannose receptor to promote their fusion.

### RNAseT2 affects actin organization in MuSCs

To investigate gene function during human MuSC fusion, we defined fusion kinetics in MuSCs. Following culture in differentiation medium, cells were fused by 20% at day 1, increasing to 30% and 50% at days 3 and 5, respectively (Fig.S2A). We showed that expression of the fusogen *MYOMAKER* appeared at day 1 and increased as the cells were fusing, while that of the fusion regulator *MYOMIXER* remained stable (Fig.S2B). Accordingly, the expression of the fusion inhibitor, the *CD9* tetraspanin, decreased with time (Fig.S2B). Based on these observations we therefore chose day 3 to investigate how RNAseT2 affects MuSC fusion.

Actin remodeling, and particularly cortical actin bundling is required for myoblast fusion ^5,18^. MuSCs treated with RNAseT2 showed a higher phalloidin intensity in their cytoplasm, indicative of changes to actin organization (Fig.2A). We further showed that RNAseT2 increased the F/G actin ratio in MuSCs (Fig.S2C), confirming a direct action of RNAseT2 on actin polymerization. Co-immunolabelling suggested that RNAseT2 partially colocalized with actin. We investigated whether colocalization of high intensity actin labeling (top 50% intensity) with high intensity RNAseT2 labeling (top 50% intensity) in MuSCs was associated with differential cell behavior.

**Figure 2.**
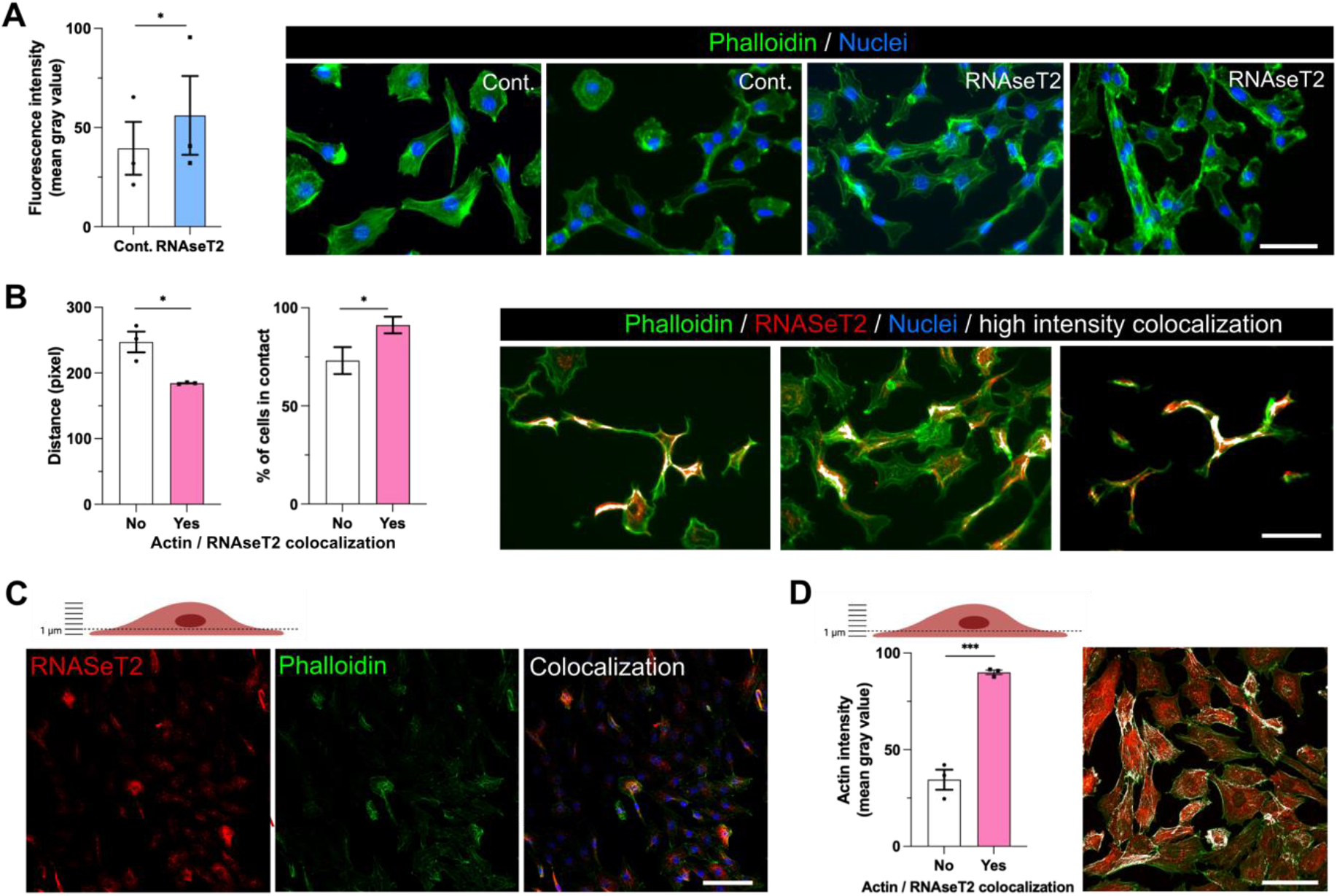
RNAseT2 induces actin remodeling in MuSCs. MuSC were cultured with or without RNAseT2 (5 µg/ml) in differentiation medium for 3 days and stained with Phalloidin-Atto488 (green) and immunostained for RNAseT2 (red) (blue = Hoechst). (**A**) Actin intensity measured in the whole cell cultures. (**B**) Distance to nearest cell (left graph) and percentage of cells in contact with another cells (right graph) was calculated for MuSCs presenting (pink bars) of not (white bars) high intensity colocalization of actin and RNAseT2 in their cytoplasm (colocalization is shown in white). (**C**) Confocal analysis of the first micron from the bottom of MuSCs showing colocalization of actin and RNAseT2. (**D**) Actin intensity was quantified from (C) in areas of colocalization with RNAseT2 (pink bar) and in the rest of the cells (white bar). Results are means ± SEM of 3 independent experiments (*p<0.05, ***p<0.001 using paired (A) and unpaired (B,D) t-test). Bars = 50 µm.

We found that cells showing actin/RNAseT2 colocalization showed increased proximity with other cells (1.35-fold closer to another cell) and showed increased contact with other cells (1.4-fold more in contact with another cell) (Fig.2B). Using confocal microscopy, we analyzed the actin and RNAseT2 distribution in the basal edge within the cells (within the first micron from the bottom) and observed a clear colocalization of RNAseT2 with actin (Fig.2C). We found that actin intensity was 2.6-fold higher in the area also labelled for RNAseT2, suggesting higher polymerization of actin when it colocalized with RNAseT2 (Fig.2D).

Altogether, these results show that RNAseT2 modulates actin remodeling towards polymerization, that RNAseT2 co-localizes with actin at sites of polymerization and this is associated with increased fusion of human MuSCs.

### RNAseT2 promotes actin reorganization through the association with the SLK STE20-like serine/threonine-protein kinase

Previous studies have shown that RNAseT2 may directly bind actin ^19,20^. To identify RNAseT2 binding partners in myoblasts, and because macrophages are difficult to transfect, we screened the C2C12 myogenic cell line using an RNAseT2-eGFP fusion protein ^21^. Co-immunoprecipitation using a GFP-TRAP revealed that RNAseT2 did not co-immunoprecipitate with actin (Fig.S3A). This indicates that RNAseT2 action on actin is indirect. To identify the binding partners of RNAseT2 in C2C12 cells, we therefore analyzed co-immunoprecipitation eluates using mass spectrometry (MS)-based quantitative proteomics. Five proteins were found significantly enriched with RNAseT2 (Table S2, Fig.S3B). Among them, SLK STE20-like serine/threonine-protein kinase was deemed an interesting candidate, since some of its functions relate to actin organization ^22,23^ and myogenic cell fusion ^24,25^.

To confirm the interaction between RNAseT2 and SLK in MuSCs, we performed Proximity Ligation Assay that permits detection of protein-protein interactions at distances lower than 40 nm. In cells treated with RNAseT2, we observed a 171% increase in the number of PLA reaction particles per cell, as well as of their fluorescence intensity (245%), confirming the high proximity between RNAseT2 and SLK in MuSCs (Fig.3A). In a functional assay, silencing of SLK (using siRNA, which decreased the SLK expression by 55% in MuSCs, Fig.S3C) blunted the effect of RNAseT2 in enhancing MuSC fusion (Fig.3B). In smooth muscle cells, SLK has been shown to indirectly activate N-WASP via its phosphorylation ^26^. N-WASP is a major actor in actin remodeling at the time of MuSC fusion ^27,28^. We therefore tested the effect of silencing SLK in MuSCs and found this resulted in a decrease in both the area (−27%) and number (−40%) of phospho-N-WASP clusters at the tips of the cells after RNAseT2 treatment relative to control RNAseT2 treated cells (Fig.3C). Eventually, in the same conditions, silencing SLK in RNAseT2 treated MuSCs decreased the area of actin filaments back to the control value (Fig.3D). SLK being shown to promote paxillin phosphorylation, resulting in actin polymerization via N-WASP phosphorylation and activation ^26^, we silenced paxillin in MuSCs. Similarly, the increase in the number of phospho-N-WASP and actin bundling area observed upon RNAseT2 treatment was blunted in the absence of paxilline (Fig.3E,F). These results show that RNAseT2 interacts with SLK to promote N-WASP phosphorylation, via paxilline, triggering actin bundling and MuSC fusion.

**Figure 3.**
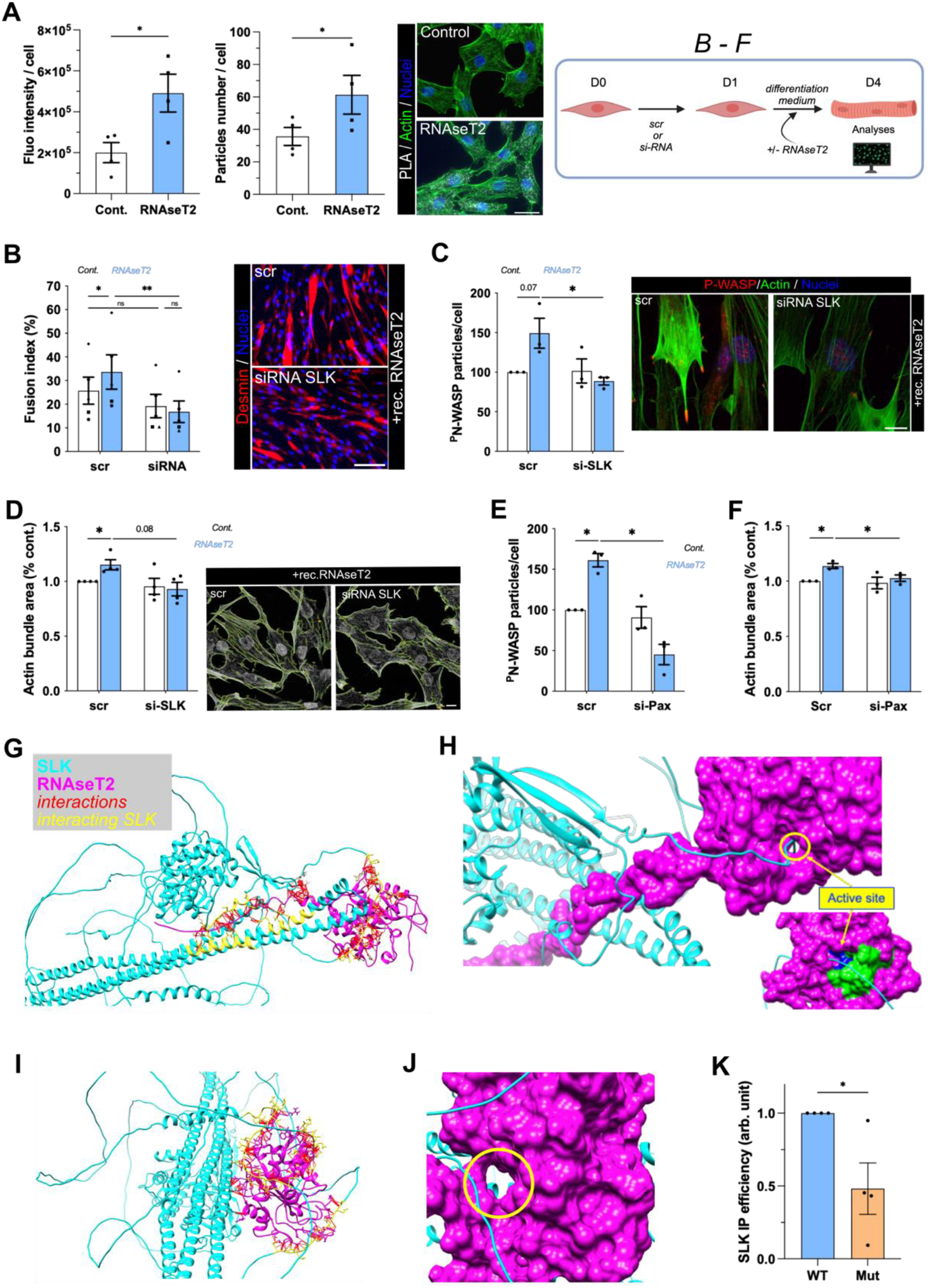
SLK-RNAseT2 interaction. **(A)** Proximity Ligation Assay in MuSCs treated or not with RNAseT2 (5 µg/ml). The intensity (left graph) and number (right graph) of the PLA particles were quantified. **(B-F)** MuSCs were transfected with scramble (Scr) or SLK-siRNA (B-D) or Paxilline-siRNA (E,F) and further incubated with or without RNAseT2 (5 µg/ml). In **(B)**, fusion was evaluated after desmin (red) and nuclei (Hoechst, blue) labeling. In **(C**,**E)**, the number of phosphorylated N-WASP puncta was quantified. In **(D**,**F)**, the area of actin filaments was quantified. **(G-J)** Model of RNAseT2-SLK interaction. **(G)** SLK (cyan) and RNAseT2 (magenta) interactions (red lines) generated using UCSF CHIMERA molecular graphic software (yellow are the portions of SLK involved in the interaction). **(H)** RNAseT2 active site exhibits a pocket in its 3D structure that accepts and interacts with a loop of the SLK protein. **(I)** Interaction (red) of mutated RNAseT2 with SLK (yellow are the portions of SLK involved in the interaction). **(J)** The mutated active site of RNAseT2 (yellow circle) does not interact with the SLK loop. **(K)** WT and mutated (Mut) RNAseT2 were immunoprecipitated from C2C12 cells and SLK presence was analyzed and quantified. Results are means ± SEM of at least 3 independent experiments (*p<0.05, **p<0.01 using paired t-test (A), unpaired t-test (K) and 2-way ANOVA (B-F)). Bars = 50 µm in A,B. 10 µm in C,D.

To explore the chemical interactions of SLK with RNAseT2, the direct docking of SLK and RNAseT2 was evaluated using the ClusPro 2.0 webserver (Fig.3G). The docking analysis, suggesting a direct interaction between RNAseT2 and SLK, revealed the occurrence of 7 salt bridges, 25 hydrogen bonds and 336 non-covalent interactions between the partners (Fig.S3D). Most identified interactions involved 3 specific regions of the SLK protein: 333-349 and 699-723 (2 regions of the protein containing polar residues), and 957-1171 (ATH Domain, previously shown to be involved in the regulation of SLK functions ^23,29^). As shown in Fig.S3E, the 3D structure analysis revealed a potential interaction of the ATH domain with the C-terminus of the RNAseT2 protein. This suggests the existence of the tight and mutliple chemical interactions between the two proteins. Moreover, the 3D structure analysis revealed that the RNAseT2 pocket (the active site) can perfectly accept and interact with a loop of the SLK protein (309-351 disordered region) (Fig.3H and S3F). The 63-69 β-strand and the 107-124 α-helix contained the residues crucial for the ribonuclease activity of RNAseT2: His65, Glu114, and His118. Strinkingly, the docking analysis suggested that mutation of His65 and His118 in RNAseT2 catalytic site, as we previously showed ^21^, could alter its interaction with SLK. Indeed, the molecular docking analysis performed between SLK and mutated RNAseT2 revealed that the regions of the SLK protein involved in the complex formation are disordered regions: 394-406, 448-471, 598-612 (Fig.3I). Most importantly, the interactions occurring at the protein-protein interface did not involve the active site of RNAseT2 in the mutant protein (Fig.3J) when compared with the WT (Fig.3H). We then immunoprecipitated mutated RNAseT2 to adress the functionality of this interaction and we observed a 50% decrease of co-immunoprecipitated SLK (Fig.3K, Fig.S3G), indicating that the interactions between the 2 proteins was lost in the mutant.

Altogether, these results suggest that RNAseT2 exerts its fusogenic activity through binding to SLK and phosphorylation/activation of N-WASP. This induces actin bundling necessary for fusion.

### Macrophage-secreted RNAseT2 stimulates MuSC fusion *in vivo*

To demonstrate that macrophage-derived RNAseT2 affects MuSC fusion *in vivo* and *ex vivo*, we evaluated its function in mouse, zebrafish and human.

To understand how MuSCs respond to secretion of RNAseT2 in macrophages *in vivo*, transgenic zebrafish over-expressing RNAseT2 under the control of a macrophage specific promoter were injured and MuSC behaviour observed using a *pax7a:egfp* transgene as previously described ^30^. Zebrafish *rnaset2* was cloned into an expression construct with a UAS driver (Fig.S4A). A stable transgenic line was established using the mpeg:gal4 driver to over-express dUAS:rnaset2_dsRed in macrophages (Fig.S4B). Transgenic animals expressing pax7a:egfp, mpeg:gal4 and dUAS:rnaset2_dsRed transgenes were injured and analyzed 4 days post-injury (dpi) as described previously ^31^. At this stage, multinucleated myofibers newly formed from GFP^pos^ MuSCs are visible. There were significantly more GFP^pos^ myofibers in the regenerating myotome of *rnaset2* over-expressing larvae (Fig.4Aa’-a’’’) as compared with control animals (Fig.4Aa-a’’) (Fig.4A left graph). We also observed a significant increase (+137%) in the ratio of myonuclei per fiber, indicative of a higher rate of MuSC fusion (Fig.4A right graph). To understand whether the elevated fusion observed at 4 dpi was associated with modified dynamics of the regenerative response, we quantified MuSCs at several stages post injury. At 1 dpi, an increased number of GFP^pos^ MuSCs within the myotome was observed in larvae expressing *rnaset2* in macrophages. Using BrdU labelling this was found to correspond to an increase in GFP^pos^ cell proliferation when compared to injured control animals (Fig.S4C). Thus, to confirm that overexpression of RNAseT2 stimulated myoblast fusion itself, we performed a temporal global over-expression of *rnaset2* using a Heat Shock inducible transgene, HS:gal4, at stages in which MuSC fusion is occuring (3 dpi). To determine how this temporal *rnaset2* expression affected fusion the myofibers and nuclei arising from *pax7a*-expressing cells were visualised 1 day after *rnaset2* over-expression using the *pax7a:egfp* transgene as it perdures after differentiation and fusion. Animals in which *rnaset2* was over-expressed showed GFP^pos^ myofibers in their regenerating myotome as expected (Fig.4Bb’). The number of regenerating GFP^pos^ myofibers was similar in control and *rnaset2* over-expressing animals (Fig.4B left graph). However, there was an elevated ratio of myonuclei (+122%) per GFP^pos^ fiber in animals over-expressing *rnaset2* from 2 dpi, indicating an increased fusion of MuSCs when *rnaset2* was expressed at later stages of regeneration (Fig.4B right graph).

**Figure 4.**
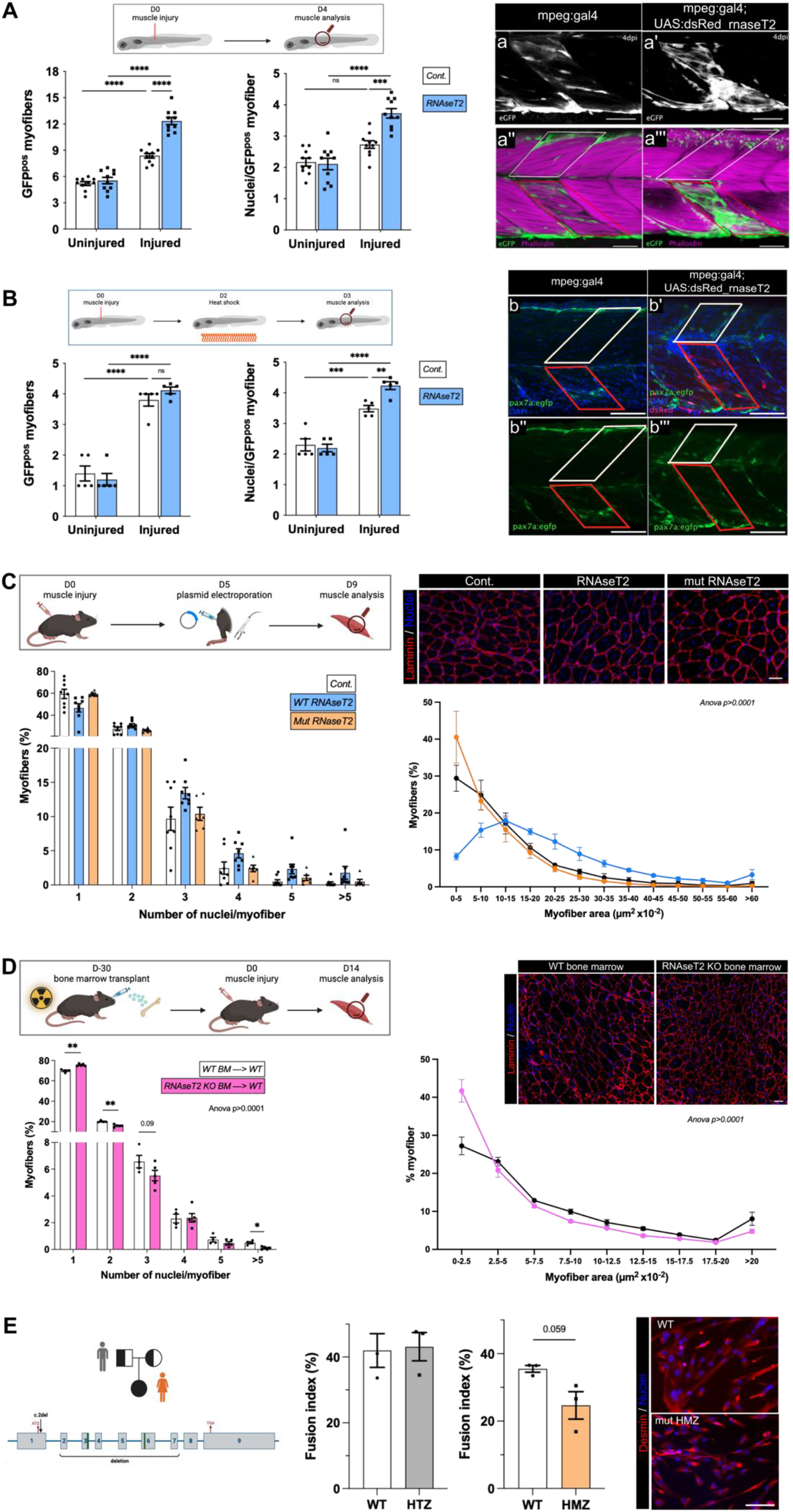
RNAseT2 stimulates muscle cell fusion *in vivo*. **(A)** Transgenic mpeg:gal4; dUAS:rnaset2_dsRed and control mpeg:gal4 zebrafish larvae, both expressing a *pax7a:egfp* transgene, were injured at 4 dpf, fixed 4 days later and labelled with phalloidin (magenta) (a’’-a’’’). The number of GFP^pos^ myofibers (left graph) and of myonuclei number per GFP^pos^ myofiber (right graph) were counted in injured (red box in a-a’’’) and uninjured (white box in a-a’’’) myotomes from the same animal. **(B)** Transgenic *hs:gal4; dUAS:rnaset2_dsRed* larvae and control *hs:gal4* larvae expressing the *pax7a:egfp* transgene were injured at 3 dpf then heat shocked at 2 days post-injury. Larvae were fixed at 24 h post heat shock (b-b’’’). The number of GFP^pos^ myofibers (left graph) and of myonuclei number per GFP^pos^ myofiber (right graph) were counted in injured (red box in b-b’’’) and uninjured (white box in b-b’’’) myotomes from the same animal. Results are means ± SEM of 5 independent experiments (**p<0.01; ***p<0.001; ****p<0.0001; not significant ns using 2-way ANOVA). Bars = 50 µm. **(C)** Mouse *tibialis anterior* muscle was electroporated with plasmid encoding for RNAseT2 or mutated RNAseT2 (or empty plasmid as a control) 5 days after injury. The number of myonuclei in regenerating myofibers (left) and the cross section area of regenerated myofibers (right) were quantified 4 days later. Results are means ± SEM of 4 independent experiments (2-way ANOVA). Bar = 50 µm. **(D)** WT mice were irradiated and transplanted with bone marrow from WT or *Rnaset2*^*-/-*^ mice. One month later, *tibialis anterior* muscle was injured and the number of myonuclei in regenerating myofibers (left) and the cross section area of regenerated myofibers (right) were quantified 14 days later. Results are means ± SEM of 4-5 experiments (2-way ANOVA). Bar = 50 µm. **(E)** Conditioned medium from macrophages issued from human patients bearing heterozygous (HTZ) and homozygous (HMZ) mutation in the *RNASET2* gene and age-matched controls was added to MuSCs for 3 days and cells were analyzed for fusion after desmin (red) and nuclei (Hoechst, blue) labelling. Results are means ± SEM of 3 independent experiments. Bar = 50 µm.

We then electroporated regenerating muscles of mice with RNAseT2; as this is a secreted molecule it will be released into the regenerating milieu of the muscle ^32^. Electroporating of RNAseT2 5 days after the injury increased the amount of RNAseT2 in the regenerating muscle (Fig.S4D). This led to an increase of the number of nuclei per myofiber 4 days later with the appearance of myofiber sections harboring 4 or more nuclei and an increase of the size of regenerating myofibers (Fig.4C). When the RNAseT2 mutant for the active site was electroporated in the same conditions, no effect was observed on the number of myonuclei per myofiber (Fig.4C). This accords with the *in vitro* results above indicating that the link with SLK is reduced in the mutant version of RNAseT2. We then performed loss of function experiments by transplanting bone marrow from *RnaseT2*^*-/-*^ mice into WT recipients. As 75% of immune cells are macrophages in the regenerating muscle, this strategy is an acceptable surrigate for investigating the role of macrophages in muscle regeneration ^33,34^. Irradiation delaying the process of regeneration ^34,35^, muscles were analyzed 14 days after injury. Deficiency of RNAseT2 in immune cells led to a decreased of the number of nuclei/myofibers, indicative of a fusion defect, associated with a decreased cross section area of the new myofibers (Fig.4D). Finally, to test whether RNAseT2 has a similar function in humans, we obtained macrophages from a human patient exhibiting a homozygous mutation in the RNAseT2 gene, triggering the deletion of the initiating codon and that of 5 exons (Fig.4E). Conditioned medium from these macrophages did not sustain the increase of MuSC fusion observed with macrophages coming from an age-matched healthy person or with macrophages bearing one mutated *RNAseT2* allele (Fig.4E). These examples from three different species confirmed that RNAseT2 delivered by restorative macrophages boost MuSC fusion during skeletal muscle regeneration.

## Discussion

Although MuSCs are very effective at regenerating muscle after injury, their *in vivo* environment is critical for their efficient function. A variety of cell types sustain the various steps of adult myogenesis ^1^. Among them, restorative macrophages promote the last steps of myogenesis including fusion ^4,6^. The molecular mechanisms underlying the profusogenic effect of restorative macrophages on MuSCs are still elusive. We have previously shown that they upregulate the expression of the metabolic regulator PPARγ which induces the expression of the growth factor GDF3 that itself directly stimulates MuSC fusion ^34^. In the present study, we show evidence for a new mechanism by which restorative macrophages stimulate MuSC fusion. They secrete the glycoprotein RNAseT2 which enters, at least in part, into MuSCs via the mannose receptor. In the cell, RNAseT2 binds to SLK, induces Paxillin phosphorylation, promoting in turn the phosphorylation of N-WASP, actin polymerization and actin bundling, which are necessary for MuSC fusion. We demonstrate *in vivo* the beneficial effects of RNAseT2 on MuSC fusion during muscle regeneration in two animal models and highlight how macrophages from human patients with *RNASET2* mutations lack this ability, highlighting its conserved function in vertebrates.

RNAseT2 is highly conserved among prokaryotes and eukaryotic species, including plants and animals. This secreted glycoprotein has been involved in a variety of cellular processes depending on its localization and the cell type involved, and was also shown to exert both structural (non-enzymatic) and enzymatic ^36^ functions. RNAseT2 has been associated with actin remodeling in a variety of cell types, and particularly subcortical actin polymerization ^37-39^. Here we show that in MuSCs, RNAseT2 does not directly link actin, but is associated with SLK. It was shown that in muscle cells, SLK expression and phosphorylation status monitor subcortical actin remodeling ^23^. The ATH domain of SLK is involved in actin reorganization ^23^ and regulates SLK function through protein binding ^29^. Here the model suggests that RNAseT2 interacts with SLK through the ATH domain in MuSCs. However, it also indicates that the active site of RNAseT2 is invoved in binding with SLK, and therefore SLK activity. Indeed, a mutated form of RNAseT2, devoid of its active site, decreased its interactions with SLK by 2-fold. In smooth muscle cells, siRNA knockdown of SLK reduces the F/G actin ratio ^26^. SLK also promotes paxillin phosphorylation in smooth muscle cells and fibroblasts ^26,40^, resulting in actin polymerization via N-WASP phosphorylation and activation ^26^. N-WASP is an important regulator of actin organization by controlling Arp2/3-dependent actin branching and is necessary for MuSC fusion ^27,28^. In skeletal muscle, SLK expression increases during MuSC differentiation; inactive SLK or SLK deletion prevents the formation of myotubes, associated with alteration of paxillin distribution ^24,25^. This suggests a mechanism by which RNAseT2 binding to SLK results in actin polymerization, necessary for myoblast fusion. We showed that SLK, via paxillin, is indeed required for RNAseT2 stimulation of N-WASP phosphorylation and of actin polymerization and bundling in human MuSCs, resulting in cell fusion. Macrophages bearing a homozygous genetic deficiency of *RNASET2*, which causes leukoencephalopathy ^41^, were not capable of stimulating MuSC fusion. *In vivo* the presence of RNAseT2, but not of the mutated form, in regenerating muscle stimulated MuSC fusion in both mouse and zebrafish. Tissue specific expression of RNAseT2 by macrophages in zebrafish resulted in a similar increase to the fusion rate, indicating that RNAseT2 released by macrophages can regulate MuSC fusion *in vivo*. Inversely, RNAseT2 deficient macrophages are defective in providing the adequate profusogenic niche to MuSCs at the time of myofiber formation.

We here report a preferential expression of RNAseT2 in restorative macrophages *in situ* and its expression directly correlated with their profusogenic properties. Previous reports have highlighted the complexity of the link between RNAseT2 and inflammation. RNAseT2 modulates the inflammatory status of macrophages *in vitro* ^14^, is involved in intracellular degradation of ssRNAs upon infection leading to TLR8 activation in macrophages ^42^, but supports dendritic cell skewing toward a Th2 profile ^16^ and promotes an inflammatory response necessary for tissue remodeling in the medicinal leech *Hirudo verbana* ^43^. The present study did not explore how RNAseT2 may affect programming of immune cells, but has explored the impact of secreted RNAseT2 on MuSC fusion.

In conclusion, this study demonstrates a new role for the highly conserved RNAseT2 that acts exogenously to regulate MuSC fusion through a cascade of phosphorylation controlling actin bundling, thus promoting tissue repair.

## Acknowledgements

This study was supported by grants from AFM-Telethon (Grant #20008 and Alliance MyoNeurALP), from the Framework Programme FP7 Endostem (under grant agreement 241440) to BC, from the Royal Society (IE161413), Muscular Dystrophy UK (UK-17GRO-PS48-0084-KNIGHT) and the BBSRC (BB/P002390/1) to RK. GJ was supported by Fondation pour la Recherche Médicale (FRM DEQ20140329495). Proteomic experiments were partially supported by Agence Nationale de la Recherche under projects ProFI (Proteomics French Infrastructure, ANR-10-INBS-08) and GRAL, a program from the Chemistry Biology Health (CBH) Graduate School of University Grenoble Alpes (ANR-17-EURE-0003). We warmly thank the GENOM’IC Facility from Institut Cochin, Paris, with a special thanks to Sébastien Jacques.

## Author’s contribution

Conceptualization: BC, JG

Methodology: AM, BC, JG, MWG, RK

Software: CG, JG

Formal analysis: AA, AM, BC, GJ, MCLB, MWG, RK, YC

Investigation: AA, CF, ELM, GJ, MWG, RM

Resources: AA, FA, OBT, OS, KT, MK

Writing - Original Draft: BC, MWG

Writing - Review & Editing: all authors

Visualization: BC, JG, MWG, RK

Supervision: BC

Project administration:BC

Funding acquisition: BC, RK

## Data availability

The transcriptomic data are deposited at GEO (GSE246884). The mass spectrometry proteomics data have been deposited to the ProteomeXchange Consortium via the PRIDE (PMID: 34723319) partner repository with the dataset identifier PXD045581 and 10.6019/PXD045581.

## Supplementary Materials

Materials and Methods

Figs. S1 to S4

Tables S1 to S3

References

